# A map of the rubisco biochemical landscape

**DOI:** 10.1101/2023.09.27.559826

**Authors:** Noam Prywes, Naiya R. Philips, Luke M. Oltrogge, Sebastian Lindner, Yi-Chin Candace Tsai, Benoit de Pins, Aidan E. Cowan, Leah J. Taylor-Kearney, Hana A. Chang, Laina N. Hall, Daniel Bellieny-Rabelo, Hunter M. Nisonoff, Rachel F. Weissman, Avi I. Flamholz, David Ding, Abhishek Y. Bhatt, Patrick M. Shih, Oliver Mueller-Cajar, Ron Milo, David F. Savage

## Abstract

Rubisco is the primary CO_2_ fixing enzyme of the biosphere yet has slow kinetics. The roles of evolution and chemical mechanism in constraining the sequence landscape of rubisco remain debated. In order to map sequence to function, we developed a massively parallel assay for rubisco using an engineered *E. coli* where enzyme function is coupled to growth. By assaying >99% of single amino acid mutants across CO_2_ concentrations, we inferred enzyme velocity and CO_2_ affinity for thousands of substitutions. We identified many highly conserved positions that tolerate mutation and rare mutations that improve CO_2_ affinity. These data suggest that non-trivial kinetic improvements are readily accessible and provide a comprehensive sequence-to-function mapping for enzyme engineering efforts.

## Introduction

Plants, algae and photosynthetic bacteria together fix ∼100 gigatons of carbon annually using <u>r</u>ib<u>u</u>lose-1,5-<u>bis</u>phosphate <u>c</u>arboxylase/<u>o</u>xygenase (rubisco), the most abundant enzyme on earth *(1)*. Rubisco catalysis, which is slow compared to many other central carbon metabolic enzymes *(2)*, is thought to limit photosynthesis under common conditions *(3)*. Rubisco is also prone to a side-reaction with oxygen leading to the hypothesis that this apparent inefficiency is in fact a careful balance of multiple biochemical tradeoffs between rate, affinity and promiscuity *(4–7)*.

Efforts to engineer improvements to rubisco have been hampered by the low throughput of obtaining accurate measurements for its parameters including catalytic rate for carboxylation (k_cat,C_, hereafter k_cat_), CO_2_ affinity (K_C_) and specificity for CO_2_ vs. O_2_ (S_C/O_). A concentrated effort across several decades has produced several hundred biochemical measurements of natural and mutant rubiscos *(4–7)*. Collection of these measurements has been biased towards plant rubiscos and the diversity of natural rubiscos remains undersampled. Library screens and rational mutations have been used in the past to increase rubisco activity. These efforts often resulted in improved expression *(8)* but rarely led to fundamental biochemical improvements *(9)*.

Protein engineering has benefited in recent years from the introduction of machine learning approaches. One goal of such efforts is to train models with labeled protein sequence-function data from high throughput functional screens *(10–15)*. Enzyme engineering with machine learning presents an additional challenge: ideally, functional data would be decomposed into individual catalytic parameters measured in high throughput either *in vitro (16)* or *in vivo (14)*.

Here, we have developed a selection assay in *E. coli* to estimate the carboxylation fitness of >99% (8760/8835) of the single amino acid mutants of the model Form II rubisco from *Rhodospirillum rubrum* (Fig. S1). Ribose phosphate isomerase was knocked out to generate *Δrpi*, a strain which only grows on glycerol when it expresses functional rubisco (Fig. S2). We then generated a barcoded library of single-amino acid mutations of the *R. rubrum* rubisco, which we assayed in high-throughput using *Δrpi*. By varying the CO_2_ concentrations of the growth environment, we were able to estimate the CO_2_ affinities 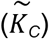 of 65% (5687) of the rubisco variants, a subset of which we went on to validate *in vitro*. This screen revealed a very small minority of mutations which improved affinity for CO_2_ ≈3-fold. These affinities have never before been observed among bacterial rubiscos, and are more typical of the Form I rubiscos found in plants and algae.

## Results and Discussion

### High-throughput characterizations of rubisco variants

The rubisco-dependent *E. coli* strain, *Δrpi*, cannot grow when glycerol is provided as the only carbon source because ribulose-5-phosphate accumulates with no outlet *(17)*. The combined actions of phosphoribulokinase (which produces the five-carbon rubisco substrate; PRK) and rubisco rescue growth by converting this otherwise dead-end metabolite into 3-phosphoglycerate, which can feed back into central carbon metabolism (Fig. 1A, S2A, for similar selection systems see *(18, 19)*).

**Figure 1:**
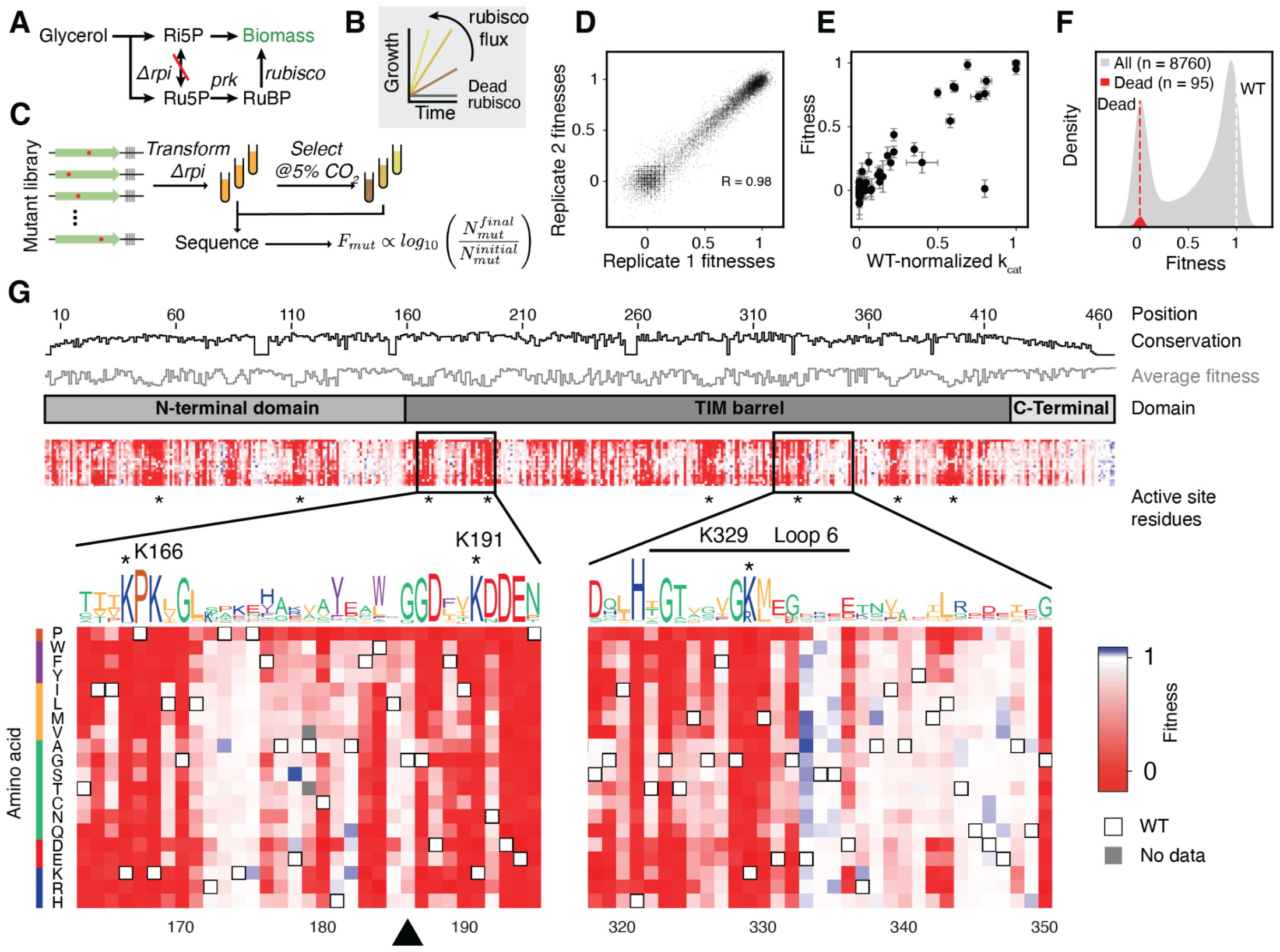
A deep mutational scan individually characterizes all single amino-acid mutations in rubisco. **A)** A summary of the metabolism of *Δrpi* the rubisco-dependent strain. **B)** *Δrpi* grows with a rate that is proportional to the flux through rubisco. **C)** Schematic of the library selection. A library of rubisco single amino-acid mutants was transformed into *Δrpi* then selected in minimal media with supplemented glycerol at elevated CO_2_. Samples were sequenced before and after selections and barcode counts were used to determine the relative fitness of each mutant. **D)** Correspondence between 2 example biological replicates, each point represents the median fitness among all barcodes for a given mutant. **E)** Fitness of 77 mutants with measurements in previous studies compared to the catalytic rates measured in those studies (k_cat_). The outlier is I190T, see supplemental text for discussion. **F)** Histogram of all variant fitnesses (grey) were normalized between values of 0 and 1 with 0 representing the average of fitnesses of mutations at a panel of known active-site positions (red distribution, average is plotted as a red dashed line) and 1 representing the average of WT barcodes (white dashed line). **G)** A heatmap of variant fitnesses. Conservation by position and the sequence logo were determined from a multiple sequence alignment of all rubiscos. Black triangle indicates G186, an example of a position with high conservation that is mutationally tolerant.

We first confirmed that the growth rate of *Δrpi* was quantitatively related to known *in vitro* enzyme behavior (Fig. 1B, S2). Expression of rubisco driven by an inducible promoter demonstrated that growth rates increased with the rubisco concentration, indicating that increased enzyme concentration led to higher fitness (Fig S2B,C, S3). Similarly, we observed faster growth in the presence of higher CO_2_ concentrations (Fig S2B,D). We next assessed whether growth-based selection correlated with biochemical behavior. Previous work on *R. rubrum* rubisco identified 77 mutants spanning <1% to 100% of wild-type catalytic rate (Supplemental Data File 1). Growth of a subset of these mutants was tested and found to correlate with reported catalytic rates (Fig. S4). Together, these results are consistent with glycerol growth of *Δrpi* being limited by rubisco carboxylation flux, which is determined by enzyme kinetics – k_cat_, K_C_ – as well as enzyme and [CO_2_] concentrations.

We next constructed a library of all single amino acid substitutions to the model Form II rubisco from *R. rubrum* (Fig. S3A). This library was cloned into a selection plasmid containing *PRK*, barcoded, and bottlenecked to ∼500,000 colonies. Long read sequencing was used to map barcodes to mutants (Fig S5B, S6) and determined that the final library contained ≈180,000 barcodes, representing 8760 mutants or >99% of the designed library (Fig S6C-F).

This library was transformed into *Δrpi* to assess mutant fitness (Fig. 1C). Mutant fitness is defined by the relative growth rate of *Δrpi* expressing that mutant. Three independent library transformations were grown in selective conditions and grown for ≈7 divisions in 5% CO_2_ (equivalent to ≈1200 μM CO_2_ in solution; wild-type K_C_ = 150 uM). Short read sequencing quantified barcode abundance before and after selection (see supplemental methods). Mutant fitness was calculated by normalizing pre- and post-selection log_10_ read-count ratios to a panel of known catalytically dead mutants and all wild-type barcodes (see Methods). Nine replicate experiments were performed with an average pairwise Pearson coefficient of 0.98 (Fig. 1D, S7).

We compared mutant fitness measurements against 77 catalytic rate values taken from the literature (Fig. 1E, Supplemental Data File 1) as well as 35 *in vitro* measurements from purified mutants (Fig. S8B) and observed a linear relationship. Overall, we observed a bimodal distribution of mutant effects (Fig. 1F) with mutant fitnesses clustering near wild-type (neutral mutations) and catalytically dead variants *(12, 20)*.

We measured fitness values for >99% (8760 out of 8835) of amino-acid substitutions (Fig. 1G, S6F, S9). Fewer than 0.14% mutations appeared more fit than WT, and when they did it was by a small amount (Fig. 1F) and 72.76% were found to be deleterious. Mutations at known active-site positions had very low fitness (e.g. K191, K166, K329, residues with asterisks Fig. 1G bottom), and mutations to proline were more deleterious on average than any other amino acid (Fig. S12). Phylogenetic conservation and average fitness at each position tended to anti-correlate (Fig. 1G top tracks, 2D, S13) consistent with previous studies *(21, 22)* however, several positions appeared to be both highly conserved and mutationally tolerant ( Fig. 1G black triangle).

### Mutational sensitivity varies across the rubisco structure

Our fitness assays revealed that some regions of the rubisco structure are much more sensitive to mutation than others (Fig. 2A.B). For example, residues on the solvent exposed faces of the structure are more tolerant to mutation, as expected, while active site and buried residues typically do not tolerate mutations well. A notable region of interest is Loop 6 of the TIM barrel, which is known to fold over the active site during substrate binding and to participate in catalysis (Fig. S1C, Fig. 2C inset). Despite this key role in catalysis, some residues in this loop are highly tolerant to mutation (e.g. E331 and E333), though the active-site residue K329 is highly sensitive (Fig. 2C).

**Figure 2:**
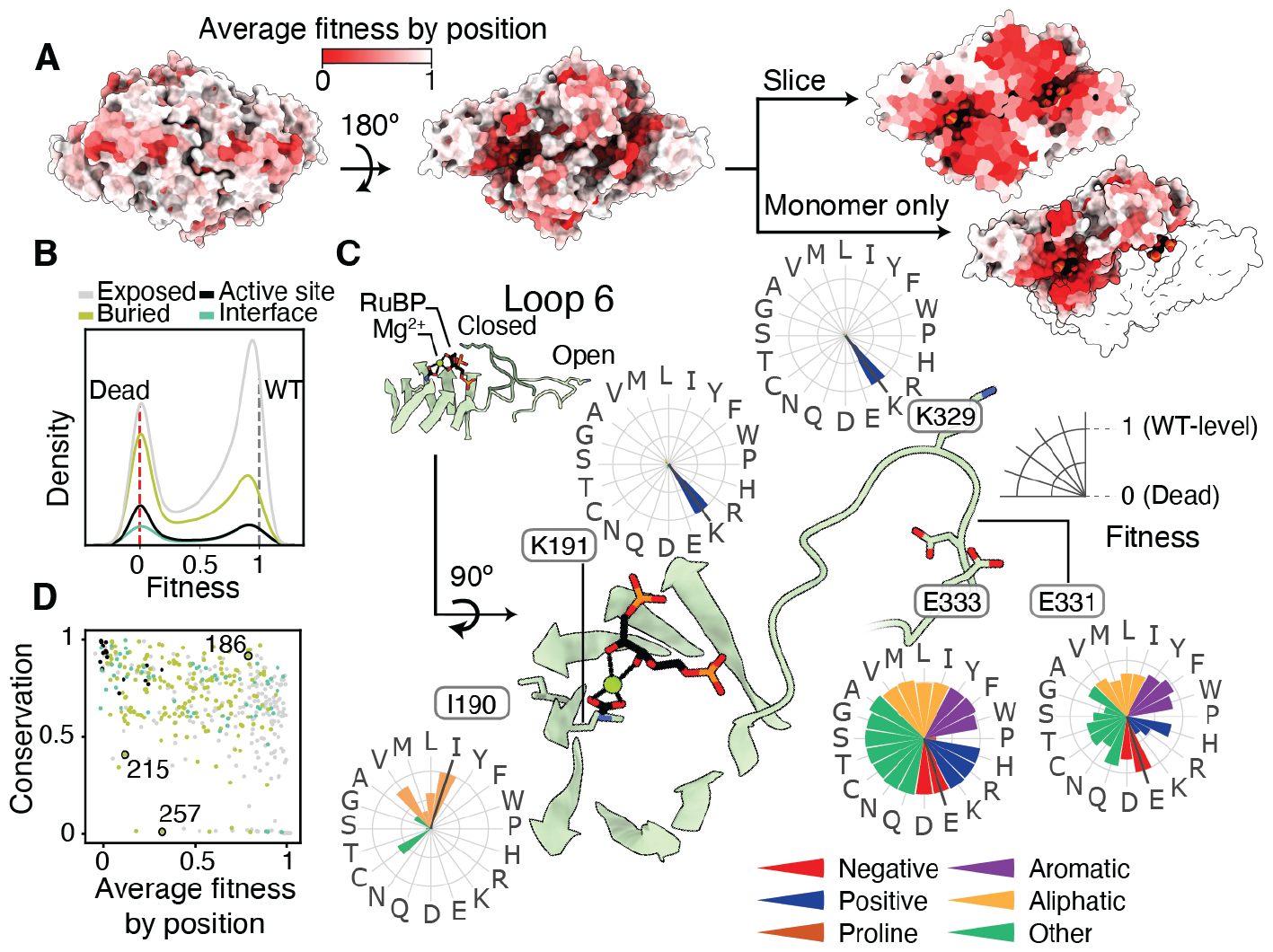
Fitness values provide structural, functional and evolutionary insights in rubisco. **A)** Structure of *R. rubrum* rubisco homodimer (Protein Data Bank (PDB) ID: 9RUB) colored by the average fitness value of a substitution at every site. **B)** Histograms of variant effects for amino-acids in different parts of the homodimer complex. **C)** Comparison of average fitness at each position against phylogenetic conservation among all rubiscos. Positions colored by the same scheme as part B. Positions 215 and 257 form a tertiary interaction, position 186 is highly conserved with no known function. **D)** Close-up view of the active-site and the mobile loop 6 region. Radar plots show the fitness effects of all mutations at a given position.

We expected that conserved positions would not tolerate mutations well. Consistent with this common hypothesis, the average fitness value at each position was negatively correlated with sequence conservation (Fig. 2C and S13). There were, however, many outliers with a number of positions being highly conserved yet showing high mutational tolerance (e.g. G186, Fig. 2D top right corner). Selection in alternative conditions may reveal what selective forces have maintained high conservation at those positions*(23)*. Positions with low conservation and low mutational tolerance may indicate a recently evolved, but critical, function *(21, 22)*; for example, M215 and H257 (Fig. 2D) are in contact in the *R. rubrum* structure but are absent in Form I sequences (Fig. S13).

### Enzyme activity and affinity can be inferred by substrate titration

Enzyme fitness is determined by the underlying biochemical parameters including catalytic rates and affinities. In order to measure these parameters individually we performed a substrate titration on the whole library of mutations in tandem (Fig. 3A). Mutant fitness values varied overall with increasing [CO_2_] (Fig. 3B, S14, S15) and some mutants’ fitnesses were strongly affected (Fig. 3C). We fit the data to a Michaelis-Menten model of catalysis to estimate effective maximum rates 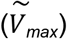 and CO_2_ half-saturation constants 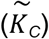 *(14)*. This fitting (Fig. 3D, see Methods) generated 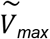 and 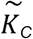 estimates for every mutant (Fig. 3G S10 and S11). We judged the reliability of the estimates by the coefficient of variation (standard deviation over the mean; *σ*/*μ)* of 1100 fits of the data for each mutation (see Methods); we focus here on the 65% of the mutants (5687) that had a coefficient of variation under 1 *(21)*. The remaining 35% are primarily mutants with low fitness values (Fig. S16) which may fail to fold altogether, though at higher expression levels or in combination with other mutations it may yet be possible to produce reliable estimates of their effects on rate and affinity.

**Figure 3:**
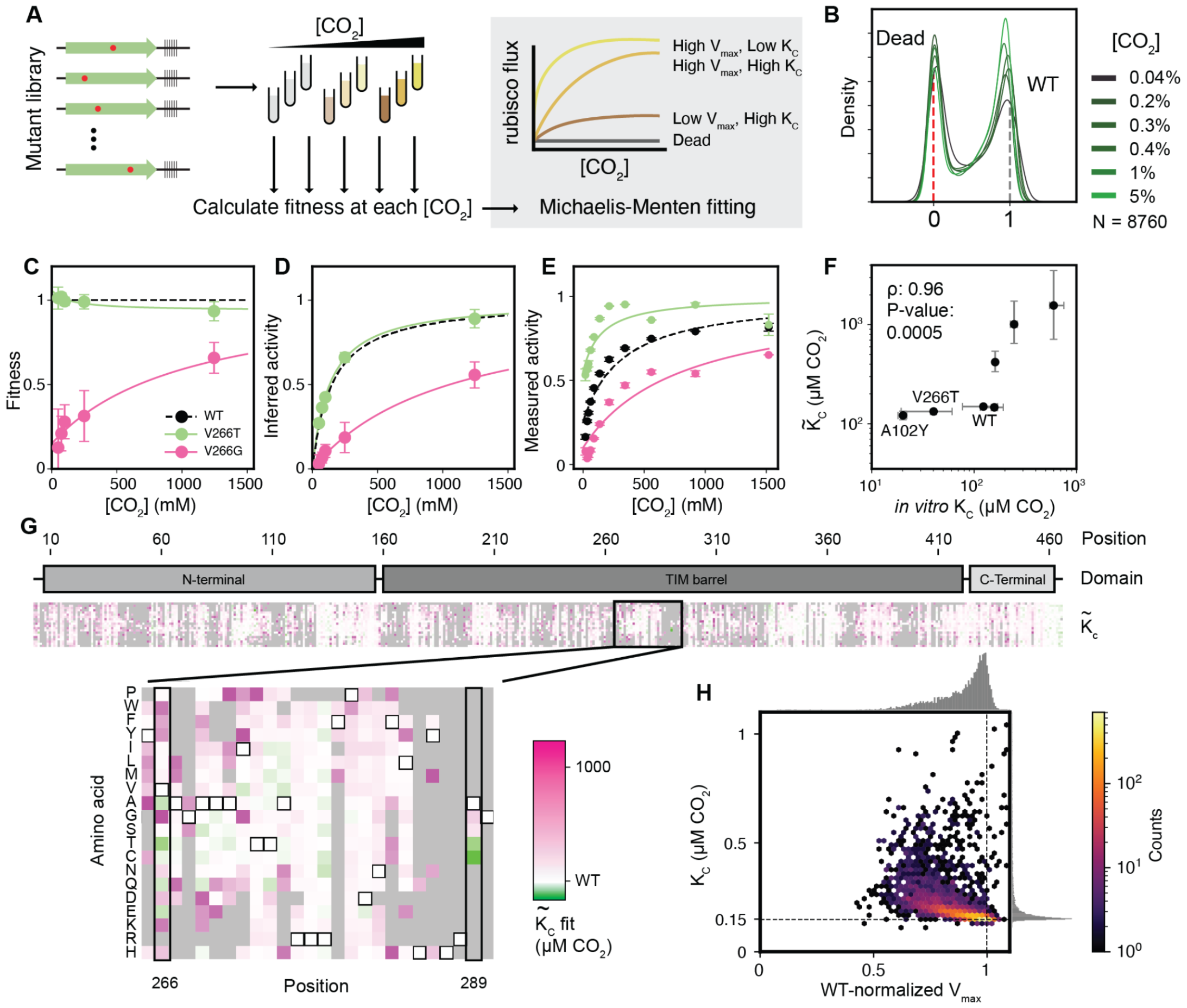
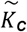 and 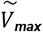 can be inferred from fitness across a CO_2_ titration. **A)** Schematic of rubisco selection in [CO_2_] titration and some examples of inferred Michaelis-Menten curves of mutants with varying K_C_ and V_Max_. **B)** Histograms of variant fitnesses at different [CO_2_]. **C)** Measured fitnesses at different [CO_2_] for two mutants. **D)** The same data as in **C** plotted under the assumptions of the Michaelis-Menten equation. **E)** Individually measured rubisco kinetics for the same two mutants from **C** and **D. F)** Comparison between *in vitro*-measured rubisco K_C_ and those inferred from fitness values 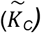. **G)** Heatmap of 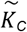 values for all mutants where the coefficient of variation is <1 (N = 5687 mutants, 65% of total). Two positions with high-affinity mutations are highlighted in the inset below. Variants where the 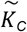 fits had a coefficient of variation above 1 are in gray. **H)** Two dimensional histogram of 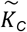 and 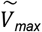 from **G** with hexagonal bins. Dashed lines represent the WT values.

We validated our 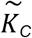 estimates by purifying a set of 7 mutants chosen to span a range of predicted 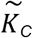 values and measuring their CO_2_ affinities *in vitro* (Fig. 3E). Unexpectedly, for several mutants, the *in vitro*-measured K_C_ values were substantially lower (i.e. tighter affinity) than expected from our prior estimates based on fitness data. For example, the 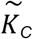 of V266T was ≈130μM but K_C_ was determined to be ≈80μM CO_2_ (Fig. 3G highlighted box, Fig. 3F).

Our estimates of 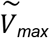 correlated with fitness (r = 0.93, Fig. S16) indicating that it is the primary driver of rubisco flux. However, V_max_ = k_cat_ x [rubisco] so variation in V_max_ can have two potential causes: rubisco expression level and 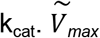 estimates report the product of those two factors.

We further found that 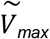 and 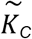 estimates anticorrelate for variants with near-WT kinetics where the estimates are most reliable (Fig. 3H). This correlation implies that, in the absence of selective pressure, the majority of single amino acid mutations harm CO_2_ affinity and V_max_ in tandem. As there is no binding site for CO_2_ in the enzyme *(24)*, this trend may be related to subtle changes in the electronics of the active site or the geometry of the bound sugar substrate before bond-formation with CO_2_. It is also possible that these effects are caused by changes to enzyme stability.

Three mutations (A289C, A102Y, V266T), caused strong improvements in CO_2_ affinity *in vivo* (Fig. 3G, 4A). Other mutations at these same positions also reduced affinity (e.g. V266G, A102F, A289G, Fig. 3C-G). These three positions are not part of the active site and sit near the C_2_ axis of the rubisco homodimer interface (Fig. 4B). In this region of the structure, residues are in closest proximity to “themselves” - i.e. to their counterpart residue in the other monomer of the homodimer. The role these amino acids play in CO_2_ entry into the active site, active site conformation, or electrostatics remains unclear.

**Figure 4:**
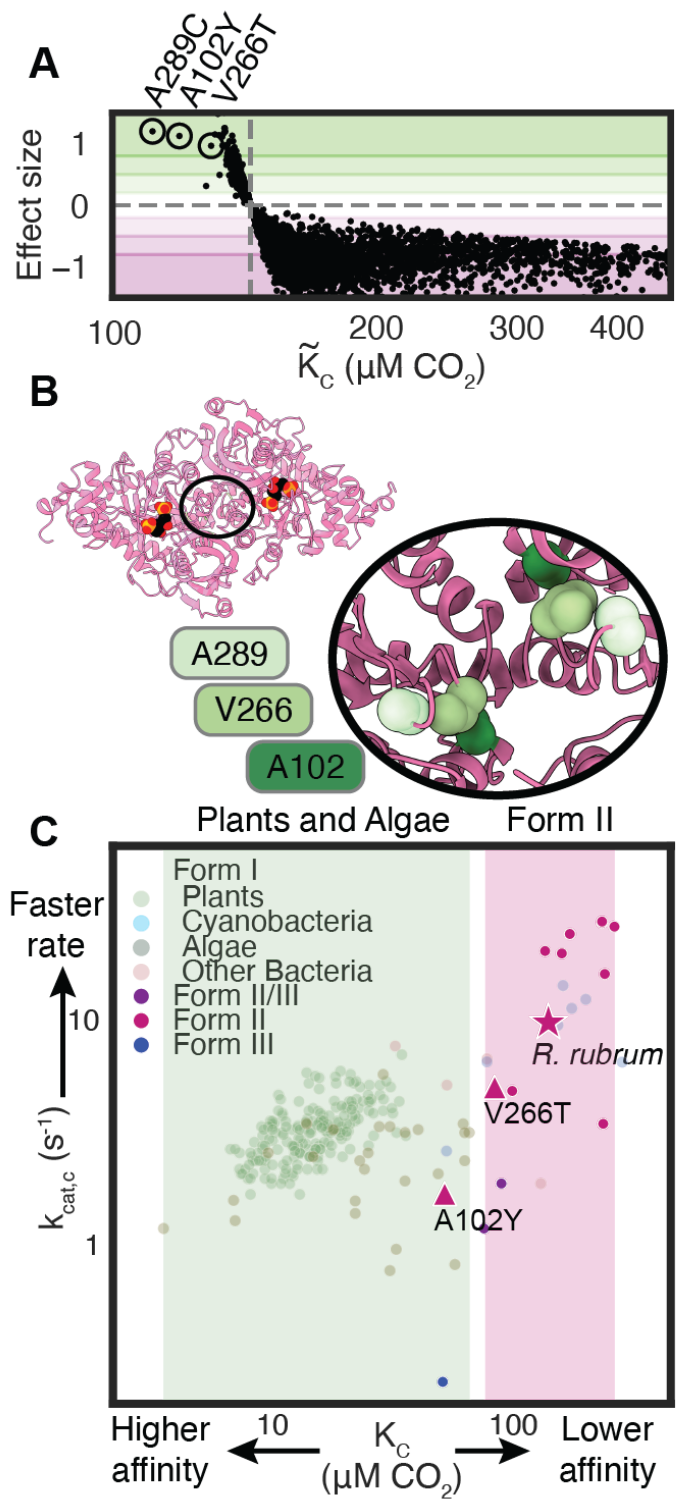
Single amino-acid mutations can traverse the functional landscape. **A)** 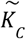 vs. effect size for each mutant. Effect size is the difference between the mutant 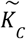 and WT K_C_ divided by the coefficient of variation of 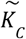. **B)** PDB structure 9RUB with inset of a zoom on the C_2_ symmetry axis. Each position appears twice due to proximity to the C_2_ axis. **C)** k_cat_ vs. K_C_ of the indicated mutants vs. all measured rubiscos from*(6, 25)*. Shaded regions indicate known ranges of 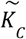 values for plants and algae in green and Form II bacterial rubiscos in pink. WT *R. rubrum* is represented by a star while mutants A102Y and V266T are triangles.

*In vitro* measurements confirmed that V266T and A102Y possess improved CO_2_ affinities (we were unable to purify A289C). This correspondence between 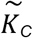 measured *in vivo* and K_C_ measured *in vitro* stands in contrast to mutations with 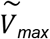 where followup biochemistry (Fig. S8B, Supplementary Data 1) did not reveal faster k_cat_ values. Variants with improved 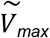 were likely improved through higher protein expression. Our 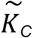 predictions were isolated from expression effects, because mutants were judged individually by their relative performance across a CO_2_ titration, and were thus more accurate. V266T and A102Y both exhibit roughly-proportional reductions in catalytic rate (Fig 4C, Table S2). The k_cat_ and K_C_ measurements place these mutants outside of the range heretofore measured among bacterial Form II variants and at the edge of the distribution of plants and algae.

## Conclusion

Among the narrow range of sequences measured here it was possible to identify mutants with substantially improved CO_2_ affinity, suggesting the enzyme parameter landscape is rugged with apparent gain-of-function readily accessible. Form I plant rubiscos typically share <50% identity with Form II bacterial rubiscos (>200 mutations, Fig S17) and are thought to have evolved under a different set of selective constraints. Furthermore, Form I and II rubiscos have different oligomeric states and Form II rubiscos lack the small subunit characteristic of Form I, so it is surprising that it is possible to traverse the functional space between them with just one amino acid change. In *R. rubrum*, the present-day sequence evolved under constraints including endogenous regulation, environmental selective pressure and possible tradeoffs between enzymatic parameters.

Various tradeoffs have been proposed in the catalytic mechanism of rubisco *(4, 6)*, including one between catalytic rate and CO_2_ affinity *(5)*. The reductions in k_cat_ observed in the mutants with the highest CO_2_ affinity is consistent with such a tradeoff (Fig. 4C). A selection of a library of higher order mutants which spans a wider range of rubisco functional possibilities could confirm or reject a tradeoff. The tradeoffs in bacterial rubiscos may also constrain the evolution of plant rubiscos. However, previous work comparing the sequence-to-function map of related proteins found substantial context-dependence on the effects of mutations *(12)*. Due to advancements in expressing plant rubiscos in *E. coli (26)*, it may be possible to use this assay to understand the biochemical constraints of the organisms which are responsible for nearly all of terrestrial photosynthesis *(27)*.

The overall space of rubiscos remains largely unexplored, raising the question of whether natural evolution has already produced rubiscos optimized for every environment. A higher throughput exploration of sequence space may reveal regions which are constrained by different tradeoffs and produce substantial engineering improvements.

## Materials and Methods

### Strains, plasmids and primers

#### Strains

Cloning was performed in a combination of *E. coli* TOP10 cells, DH5α and NEB Turbo cells. Protein expression was carried out using BL21(DE3). *Δrpi* was previously produced from the BW25113 strain by knocking out *rpiA* from the Keio strain lacking *rpiB* as well as the *edd* gene. The latter deletion makes the strain rubisco-dependent when grown on gluconate, a feature we did not make use of in this study.

#### Plasmids

Sequences and further details about plasmids used in this study can be found in supplemental data file S3.

#### pUC19_rbcL

The rubisco mutant library was assembled in a standard pUC19 vector. This plasmid was used as a PCR template for each of the 11 sublibrary ligation destination sites.

#### NP-11-64-1

Selections were conducted using a plasmid designed for this study with a p15 origin, chloramphenicol resistance, LacI controlling rubisco expression, TetR controlling PRK expression and a barcode.

#### NP-11-63

Protein overexpression in BL21(DE3) cells was conducted using pET28 with a SUMO domain upstream of the expressed gene *(25)*. pSF1389 is the plasmid that expresses the necessary SUMOlase, bdSENP1, from *Brachypodium distachyon*.

#### Primers

All primers were purchased from IDT and the oligo pool was purchased from TWIST. For sequences see supplemental data file S3.

### Library design and construction

The *R. rubrum* rubisco sequence was codon-optimized for *E. coli* and systematically mutated via the scheme outlined in Fig. S5. The rubisco gene was split into 11 pieces. For each of those pieces (≈200 bp each) all point mutants were designed and synthesized as oligonucleotide pools. 11 oligo sub-library pools, containing all single mutants within their respective ≈200 bp region, were purchased from Twist Bioscience and each sub-library was amplified individually using Kapa Hifi polymerase with a cycle number of 15. Each rubisco gene fragment was inserted into a corresponding linearized pUC19 destination vector, containing the remainder of the rubisco sequence flanking the insert, via golden gate assembly. This assembly generated 11 sub-libraries of the full-length *R. rubrum* rubisco gene with each sub-library containing a ≈200 bp region including all single mutants. Each of these 11 rubisco libraries were separately transformed into *E. coli* TOP10 cells and in each case >10,000 transformants were scraped from agar plates to ensure oversampling of the ≈1,000 variants in each sublibrary. Plasmids were purified from each sub-library and mixed together at equal molar ratios to generate the full protein sequence library.

In order to produce the final library for assay, a selection plasmid containing an induction system for rubisco and PRK (Tac- and Tet-inducible, respectively) was amplified with primers that included a random 30 nucleotide barcode. The linearized plasmid amplicon and the library were cut with BsaI and BsmBI, respectively, ligated together and transformed into TOP10 cells. Plasmid was purified by scraping ≈500,000 colonies and transformed in triplicate into *Δrpi* cells. These transformations were grown in 2XYT media into log phase (OD = 0.6) and frozen as 25% glycerol stocks.

### Long-read sequencing analysis

The plasmid library was cut with SacII and sent for Sequel II PacBio sequencing. Reads were aligned and grouped by their barcodes. All reads of a given barcode were aligned and a consensus sequence was obtained using SAMtools*(28)*. Consensus sequences were retained if they were WT or had one mutation that matched the designed library. Any mutation in the backbone invalidated a barcode. A lookup table was generated to link each barcode to its associated mutation. The *in silico* procedures described in this study are publicly available at https://github.com/SavageLab/rubiscodms.

### Library characterization and screening

Selections were performed by diluting 200 uL of glycerol stock with OD of ≈0.25 into 5 mL of M9 minimal media with added chloramphenicol (25 μg/mL), glycerol (0.4%), 20 μM IPTG and 20 nM anhydrotetracycline. These cultures were grown at 37 °C in different CO_2_ concentrations until they reached an OD at 5 mL of 1.2 +/- 0.2. This corresponds to a 100-fold expansion of the cells, i.e. between 6 and 7 doublings.

Cultures before and after selection were spun down and we lysed the cells and performed a standard plasmid extraction protocol using QIAprep Spin Miniprep Kit (QIAGEN, Hilden, Germany). Illumina amplicons were generated by PCR of the barcode region. These amplicons were sequences using a NextSeq™ P3 kit

### Calculation of variant enrichment

Variant enrichments were computed from the log ratio of barcode read counts. The enrichment calculations include two processing parameters: a minimum count threshold (c_min_) and a pseudocount constant (α_p_). The count threshold is the minimum number of barcode reads that must be observed either pre- or post-selection for the barcode to be included in the enrichment calculation. The pseudocount constant is used to add a small positive value to each barcode count to circumvent division by zero errors. We use a pseudocount value that is weighted by the total number of reads in each condition. For the *j*^th^ variant and the individual barcodes, *i*, passing the threshold condition the variant enrichment is calculated as,

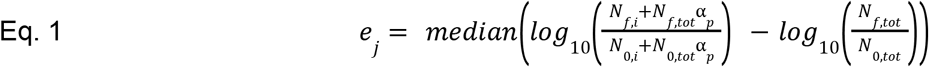

To identify optimal values for these parameters, we computed the variant enrichments across a 2D parameter sweep of c_min_ and α_p_ to find the combination that resulted in the maximum mean Pearson correlation coefficient across all replicates at each condition. These were c_min_ = 5 and α_p_ = 3.65e-7 (average of 0.3 pseudocounts) leading to a correlation coefficient of 0.978. Variant enrichment, *e*_*j*_, was then calculated for every mutant using Eq. 1.

The variant enrichments were then normalized such that wild-type has an enrichment value of 1 in all conditions and catalytically dead mutants have a median enrichment of 0. For the “dead” variant enrichment we computed the median enrichment for all mutations at the catalytic positions K191, K166, K329, D193, E194, and H287. The normalized enrichments at each condition were computed as,

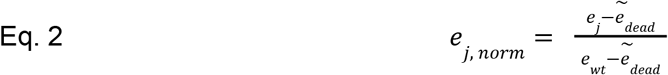

where e_j_ is the enrichment of the j^th^ variant as given in Eq. 1, e_wt_ is the wild-type enrichment, and 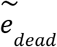 is the median enrichment across all mutants of the catalytic residues listed above.

### Michaelis Menten fits to enrichment data

The DMS library enrichments across different CO_2_ concentrations were used to estimate Michaelis-Menten kinetic parameters for every variant. Guided by the linear relationship between growth rate and k_cat_ observed in Fig. 1D we assume that the cell growth rate is proportional to the rubisco enzyme velocity to derive the CO_2_ titration fits (see SI, Derivation of Michaelis-Menten Fit).

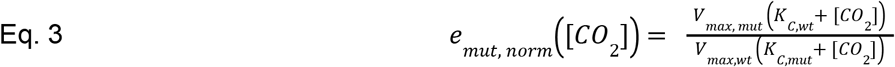

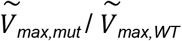 is the ratio of mutant maximum velocity relative to wild-type, 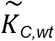 is the wild-type K_C_ for which we used the value 149 μM, and 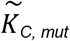 is the mutant K_C_. The titration curves in triplicate for each variant were fit to Eq. 3 using non-linear least squares curve fitting while requiring both V_max_ and *K*_*C, mut*_ to be positive.

We noted that the 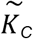 fits to certain variants – particularly ones with low 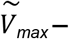 were sensitive to the choice of processing parameters c_min_ and α_p_. Given the semi-arbitrary nature of these parameters, this is clearly an undesirable dependence and engenders low confidence in the inferred 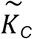 values. To account for this uncertainty we conducted a parameter sweep (with 11 different c_min_ values linearly spaced between 0 and 50, and 10 α_p_ values log spaced between 1e-9 and 1e-6), and computed the variant enrichments for all combinations of these parameters. Then we performed 10 bootstrap subsamplings of the replicates for all parameter sets and performed the ratiometric Michaelis-Menten fit.From this set of 1100 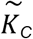 fit values for each variant we computed a quartile-based coefficient of variation that was used as a figure of merit for the 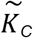.

### Multiple sequence alignment

An MSA of the broader rubisco family beyond Form II rubiscos was created using the profile HMM homology search tool jackhmmer *(29)*. Starting with the *R. rubrum* rubisco sequence, jackhmmer applied five search iterations with a bit score threshold of 0.5 bits/residue against the UniRef100 database of non-redundant protein sequences *(30)*. To compute phylogenetic conservation at each position, for each possible amino acid we computed the fraction of the total sequences that had that amino acid at the corresponding position of the MSA. The phylogenetic conservation is the maximum fraction, where the maximum is taken over all possible amino acids. Thus, if a position has an alanine in 90% of the sequences of the MSA, the phylogenetic conservation will be 0.9.

### Protein purification

*E. coli* BL21(DE3) cells were transformed with pET28 (encoding the desired rubisco with a 14x His and SUMO affinity tag) and pGro plasmids. Colonies were grown at 37°C in 100mL of 2XYT media under Kanamycin selection (50 μg/ml) to an OD of 0.3-1. 1 mM arabinose was added to each culture and then incubated at 16°C for 30 minutes. Protein expression was induced with isopropyl-b-D-thiogalactopyranoside (IPTG, Millipore) at 100 uM and cells were grown overnight at 16°C. Cultures were spun down (15 min; 4,000 g; 4°C) and purified as reported *(25)*. Briefly, cultures were spun down and lysed using BPER-II™. Lysates were centrifuged to remove insoluble fraction. His-tag purification using Ni-NTA resin (Thermo Fisher, Massachusetts, United States) was performed and rubisco was eluted by SUMO tag cleavage with bdSUMO protease (as produced in Davidi et al. 2020). Purified proteins were concentrated and stored at 4 °C until kinetic measurement (within 24 hr). Samples were run on an SDS-PAGE gel to ensure purity.

### Rubisco spectrophotometric assay

Both k_cat_ and K_C_ measurements use the same coupled-enzyme mixture wherein the phosphorylation and subsequent reduction of 1,3-bisphosphoglycerate, the product of RuBP carboxylation, was coupled to NADH oxidation which can be followed through 340 nm absorbance. Following *(31)* and *(25)* the reaction mixture (Table S1) contains buffer at pH 8, MgCl_2_, DTT, 2 mM ATP, 10 mM creatine phosphate, 0.5mM NADH, 1mM EDTA and 20U/mL each of PGK, GAPDH and creatine phosphokinase. Reaction volumes are 150 μL and samples are shaken once before absorbance measurements begin. Absorbance measurements are collected on a SPARK plate reader with O_2_ and CO_2_ control (TECAN). The extinction coefficient of NADH in the plate reader was determined through a standard curve of NADH solutions of known concentration (determined by a genesys20 spectrophotometer with a standard 1 cm pathlength, Thermo Fisher). Absorbance over time gives a rate of NADH oxidation and therefore a carboxylation rate. Because rubisco produces 2 molecules of 3-phosphoglycerate for every carboxylation reaction we assume a 2:1 ratio of NADH oxidation rate to carboxylation rate.

**Table S2.** Assay mix composition.

| Component | Assay concentration | Source |
| --- | --- | --- |
| EPPS buffer pH 8.0 | 100 mM | Alfa Aesar (Cat # J61296) |
| $MgCl_2$ | 20 mM | Sigma Aldrich (Cat # M2670-500G) |
| Dithiothreitol | 0.5 mM | Bio Basic Canada inc. (Cat # DB0058) |
| ATP | 2 mM | Sigma Aldrich (Cat # A3377-5G) |
| Phosphocreatine | 10 mM | Sigma Aldrich (Cat # 27920-5G) |
| NADH | 1.7 mM | Merck (Cat # 481913-1GM) |
| Carbonic anhydrase | 0.1 mg/mL | Sigma Aldrich (Cat # C3934-100MG) |
| Creatine phosphokinase | 20 U/mL | Sigma Aldrich (Cat # C3755-35KU) |
| Glyceraldehyde 3-phosphate dehydrogenase | 20 U/mL | Sigma Aldrich (Cat # G2267-10KU) |
| 3-Phosphoglyceric phosphokinase | 20 U/mL | Sigma Aldrich (Cat # P7634-5KU) |

### Spectrophotometric measurements of k_cat_

The carboxylation rate (k_cat_) of each rubisco was measured using methods established previously *(25)*. Briefly, rubisco was activated by incubation for 15 minutes at room temperature with CO_2_ (4%) and O_2_ (0.4%) and added (final concentration of 80 nM) to aliquots of appropriately-diluted assay mix (see Table S2) containing different CABP concentrations pre-equilibrated in a plate reader (Infinite® 200 PRO; TECAN) at 30°C, under the same gas concentrations. After 15 min, RuBP (final concentration of 1 mM) was added to the reaction mix and the absorbance at 340 nm was measured to quantify the carboxylation rates. A linear regression model was used to plot reaction rates as a function of CABP concentration. The k_cat_ was calculated by dividing y-intercept (reaction rates) by x-intercept (concentration of active sites). Protein was purified in triplicate for k_cat_ determination.

### Spectrophotometric measurements of K_C_

Purified rubisco mutants were activated (40 mM bicarbonate and 20 mM MgCl2) and added to a 96-well plate along with assay mix (Table S2, in this case the same concentration of Hepes pH 8 buffer was used but EPPS can be substituted). Bicarbonate was added for a range of concentrations (1.5, 2.5, 4.2, 7, 11.6, 19.4, 32.4, 54, 90 and 150mM). Plates and RuBP were pre-equilibrated at 0.3% O_2_ and 0% CO_2_ at room temperature. RuBP was added to a final concentration of 1.25 mM with water serving as a control for each replicate. NADH oxidation was measured by A340 as in the kcat assay. Absorbance curves were analyzed using a custom script to perform a hyper-parameter search to choose a square in which to take the slope as carboxylation rate that best represented the majority of the monotonic decrease in A_340_. K_C_ was derived by fitting the Michaelis-Menten curve using a non-linear least squares method. Error bars were determined depending on replicates: (1) Multi day replicates: Michaelis-Menten fits were made for each replicate, std error and median was calculated based on these fits (2) Triplicates: Absorbance data was fit 100 times using different hyperparameters. Michaelis-Menten fits of each set of rates were calculated and the median K_C_ value was plotted. Error values were determined from the K_C_ values of the hyperparameters one standard deviation above and below the median. Standard deviations and medians were calculated based on technical replicates. Subsequently, three different fits were made: one based on the median, one based on the lowest reaction rate and one based on the highest reaction rate for each point.

### Radiometric measurements of K_C_ and k_cat_

^14^CO_2_ fixation assays were conducted as in (Davidi et al. *(25)*) with minor modifications. Assay buffer (100 mM EPPES-NaOH pH 8, 20 mM MgCl_2_, 1 mM EDTA) was sparged with N_2_ gas. Rubisco, purified as described above, was diluted to ∼10 μM (quantified using UV absorbance) in assay buffer. It was then diluted with one volume of assay buffer containing 40 mM NaH^14^CO_3_ to activate. 0.5 mL reactions were conducted at 25°C in 7.7 ml septum-capped glass scintillation vials (Perkin-Elmer) with 100 μg/mL carbonic anhydrase, 1 mM RuBP and NaH^14^CO_3_ concentrations ranging from 0.4 to 17 mM (which corresponds to 15 to 215 μM CO_2_). The assay was initiated by the addition of a 20 μL aliquot of activated rubisco and stopped after 2 minutes by the addition of 200 μL 50% (v/v) formic acid.

The specific activity of ^14^CO_2_ was measured by performing a 1 hour assay at the highest ^14^CO_2_ concentration containing 10 nmoles of RuBP. Reactions were dried on a heat block, resuspended in 1 mL water and mixed with 3 mL Ultima Gold XR scintillant for quantification with a Hidex scintillation counter.

The rubisco active-site concentration used in each assay was quantified in duplicate by a [^14^C]-2-CABP binding assay. 10 μL of the ∼10 μM rubisco solution was activated in assay buffer containing 40 mM cold NaHCO_3_ (final volume 100 μl) for at least ten minutes.1.5 μL of 1.8 mM ^14^C-carboxypentitol bisphosphate was added and incubated for at least one hour at 25°C. [^14^C]-2-CABP bound rubisco was separated from free [^14^C]-2-CPBP by size exclusion chromatography (Sephadex G-50 Fine, gE Healthcare)) and quantified by scintillation counting.

The data was fit to the Michaelis-Menten equation using the concatenated data of 3-4 experiments performed on different days.

### Quantification of soluble enzyme concentration via Immunoblot

*Δrpi* strain with WT rubisco was grown under selective conditions (overnight at 37 °C in M9 media with 0.4% glycerol and 20 nM aTc) with varying IPTG concentrations at 5% CO2 for 24h. Afterwards, turbid cultures were spun down (10 min; 4,000 g; 4°C) culminating in roughly 20 mg pellet per sample. Pellets were lysed with 200 μL of BPER II and supernatant was transferred into a fresh tube and mixed with SDS loading dye. BioRad RTA Transfer Kit for Transblot Turbo Low Fluorescence PVDF was used in combination with the Trans-Blot® Turbo™ Transfer System. Nitrocellulose Membrane was carefully cut between 50 and 70 kDa post-blocking using a razor blade. Primary Anti-RbcL II Rubisco large subunit Form II Antibody from Agrisera (1:10000) and DnaK Antibody from Abcam (1:5000) were incubated separately. Secondary HRP-conjugated antibodies Donkey anti-mouse for DnaK (Santa Cruz Biotechnology) and Goat pAB to RB IgG HRP (Abcam were both used at 1:10000. Subsequently BioRad Clarity Max Western ECL Substrates were applied and the final results were imaged using a GelDoc.

## Supporting information

rbcL_DMS_supp

## Acknowledgements

We thank Niv Antonovsky and Arren Bar-Even for taking part in formulating the basis for this work as well as Naama Tepper and Shira Amram for originally conceiving of and producing the *Δrpi* strain respectively. We thank Philip Romero, Nat Thompson, Leon Fedotov, Orren Saltzman, Eden Prywes, Stacia Wyman, Bin Yu and Jack Desmarais for essential help in the process of data analysis. For their assistance in the process of generating and validating the DMS library we thank Andrew Glazer, Kenneth Matreyek, Jesse Bloom and Kim Reynolds. Additionally we thank Julia Tartaglia for the use of her sequencing primers and Netra Krishnappa for assistance in running NGS samples. We would like to thank Elaine Meng for assistance using ChimeraX. Finally we thank Flora Wang for technical assistance over the weekends.

## Funding

National Institutes of Health grant K99GM141455-01 (NP)

DFS is an Investigator of the Howard Hughes Medical Institute

U. S. Department of Energy, Physical Biosciences Program, Award Number DE-SC0016240 (DFS)

## Author contributions

Conceptualization: NP, AIF, DFS

Methodology: NP, NRP, LMO, SL, DD, OMC, RM, DFS

Investigation: NP, NRP, SL, YCT, BdP, AEC, LJTK, HAC, LNH, DBR, HMN, RFW, AYB

Visualization: NP, LMO, SL, DFS

Funding acquisition: NP, DFS

Project administration: NP, DFS

Supervision: NP, PMS, OMC, RM, DFS

Writing – original draft: NP, LNH, AIF, DFS

## Data and materials availability

All data are available in the main text or the supplementary materials.

