## Supplementary material for "A map of the rubisco biochemical landscape": rbcL_DMS_supp

### Supplement

#### Figures

##### *Rubisco structure*

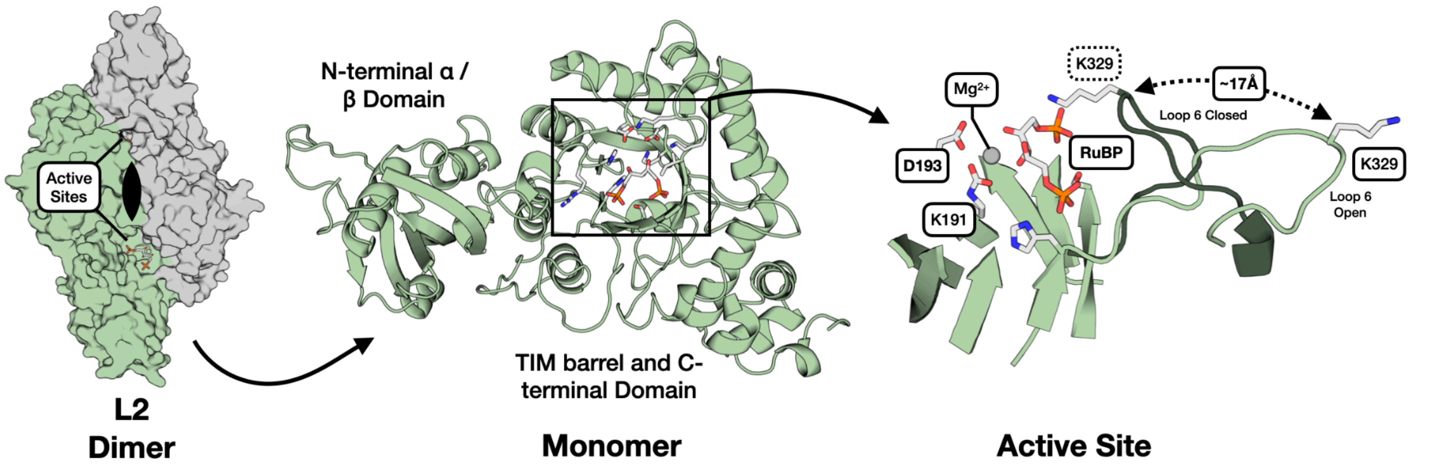


**Figure S1: *R. rubrum* rubisco structure. Left:** Overall structure of the 2-large subunit (L2) homodimer with active sites and $C_2$-symmetry axis labeled. (PDB: 9RUB). **Center:** Ribbon diagram of one monomer with the 3 subdomains labeled. View is of the interfacial side. **Right:** Close-up view of the active site. Closed form of loop 6 is from the 8RUC structure. Active site residues and RuBP substrate are labeled.

##### *Growth rate heatmap by condition*

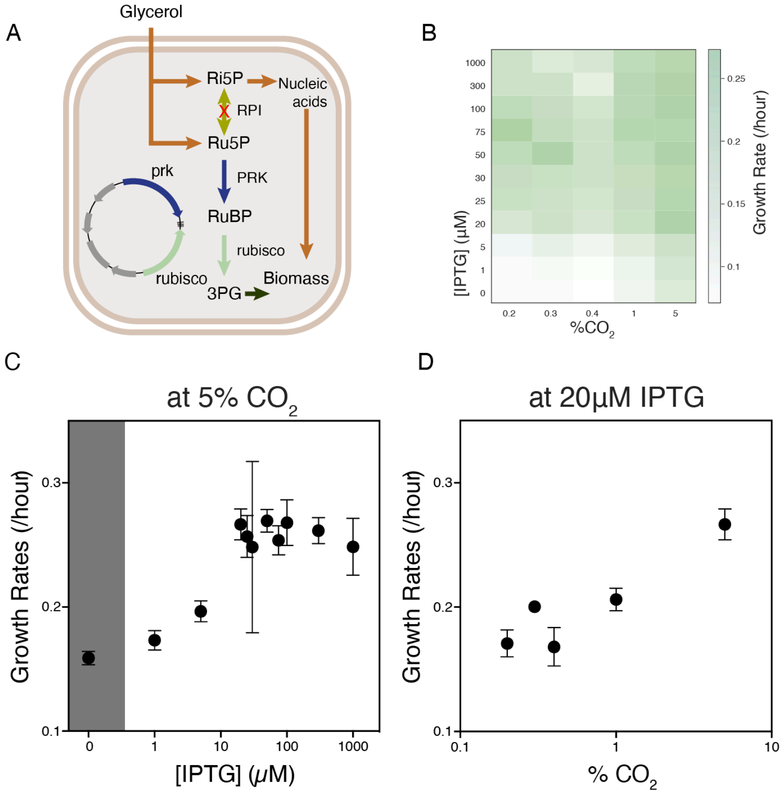

**Figure S2: *Δrpi* is a rubisco-dependent *E. coli* strain with a growth rate that correlates to rubisco flux. A)** Schematic of the *Δrpi* strain of rubisco-dependent *E. coli*. PRK and rubisco compensate for the deletion of RPI and rescue growth. **B)** A heatmap of growth rates across a two-dimensional titration of $CO_2$ and IPTG. **C,D)** Growth rates across one-dimensional titrations of rubisco induction by IPTG (**C**) and $CO_2$ (**D**) concentrations.

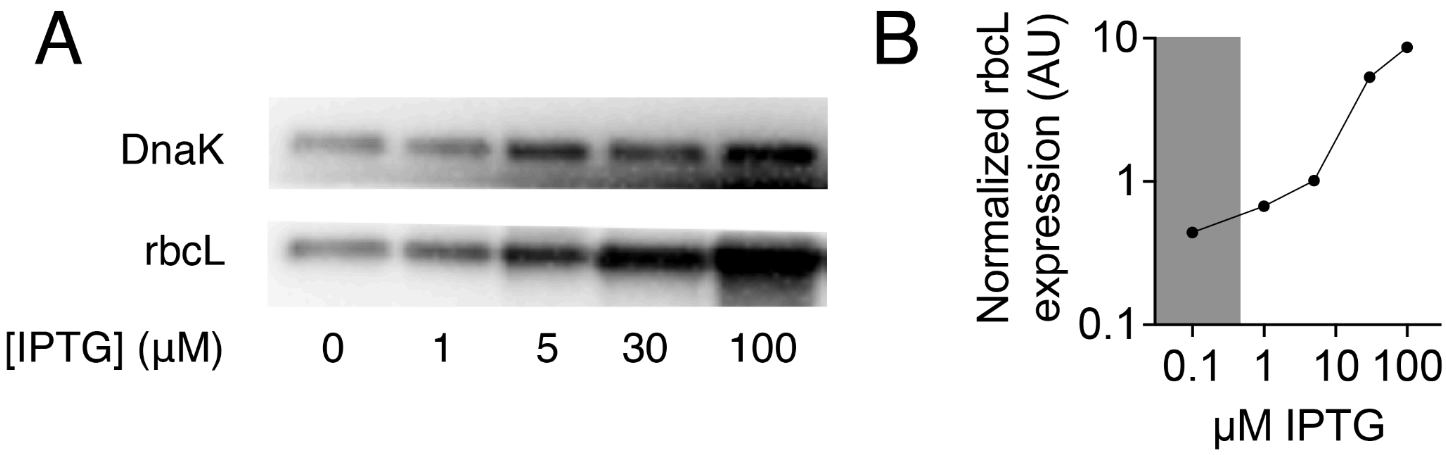


**Figure S3: Increased IPTG concentration leads to higher rubisco expression. A)** Immunoblots for soluble rubisco with DnaK as a loading control. Samples are of *Δrpi* cells grown in selection media (see Methods) with different concentrations of IPTG. **B)** Ratio of band intensities as a function of IPTG concentration.

*Mutant comparisons*

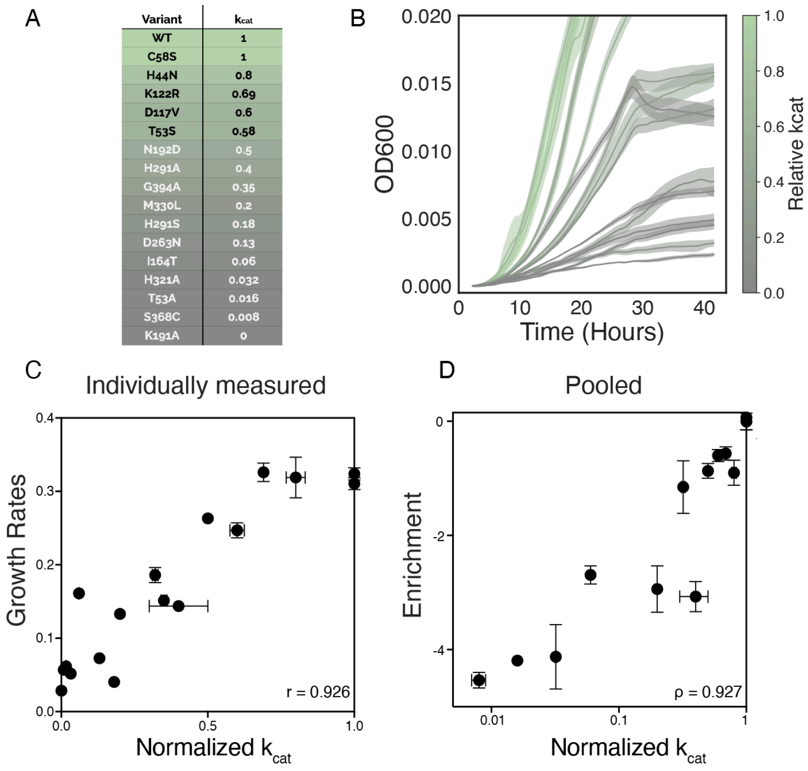


**Figure S4: *Δrpi* grows with a rate proportional to mutant $k_{cat}$: A)** A panel of mutants from the literature and their associated $k_{cat}$ measurements normalized to WT. The WT value is ≈11/s. **B)** Growth curves of *Δrpi* expressing the mutants from A. Coloring in A and B is on the same scale and reflects $k_{cat}$ values from the literature. **C)** Growth rates calculated from the curves in **B**, plotted against the normalized $k_{cat}$ values. **D)** Mutant enrichments for the same mutants as in **C** measured in one nanopore sequencing experiment. Error bars in **C** determined as standard deviations of three or more replicates. Error bars in **C** determined as standard deviations of three different barcodes for each mutant. Errors in literature values are shown from studies where they were reported.

#### *Library construction and analysis*

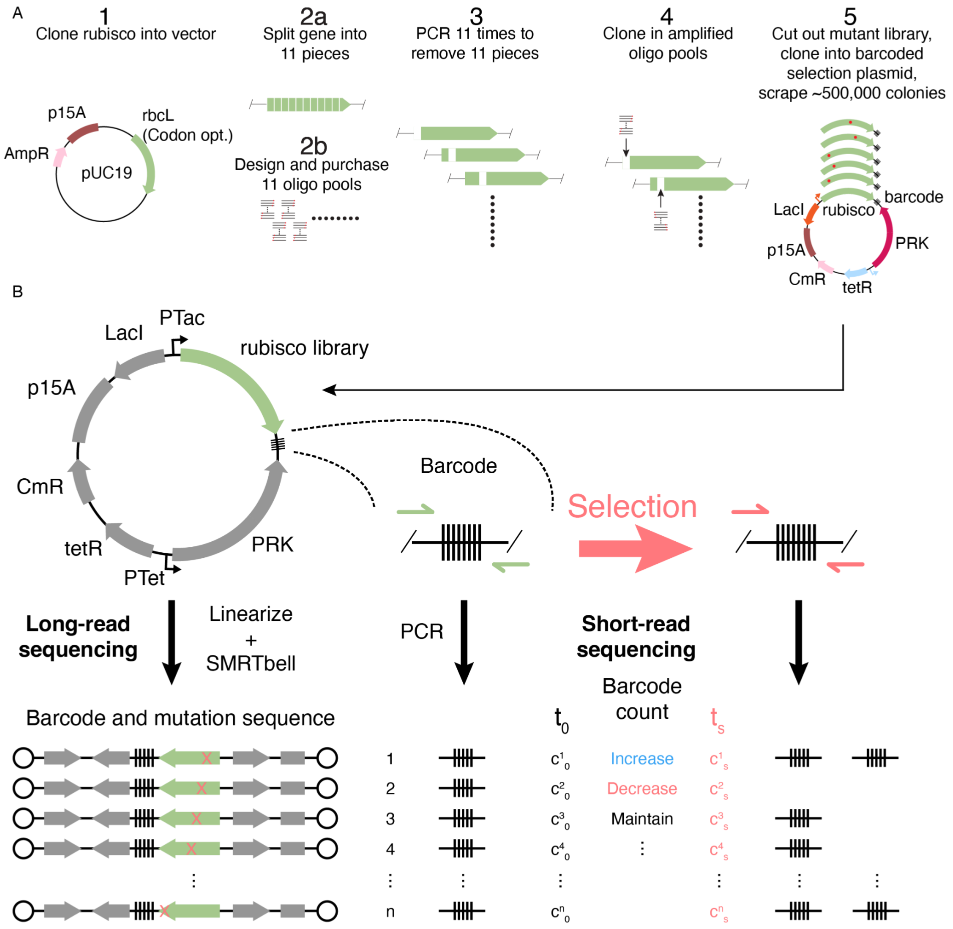


**Figure S5: Library construction and characterization pipeline. A)** Library construction procedure. **Step 1)** Clone a codon-optimized *R. rubrum* rubisco sequence into pUC19. **Step 2a)** Choose locations to split the gene which are appropriate for the cloning of subpool libraries. **Step 2b)** PCR amplify the sublibraries from an oligo pool containing all 8778 mutations. **Step 3)** PCR amplify the backbone with a space missing for the ligation of an oligo subpool. **Step 4)** Ligate each oligo subpool to its appropriate backbone. **Step 5)** Combine the sublibraries, cut the full, mutated genes out and ligate them into a PCR-amplified and barcoded backbone. After transformation scrape the desired number of colonies for selection. **B)** Library sequencing strategy. The library was characterized by long read sequencing. Barcode abundances were measure by short-read sequencing before and after selection (see methods).

#### *Library characterization*

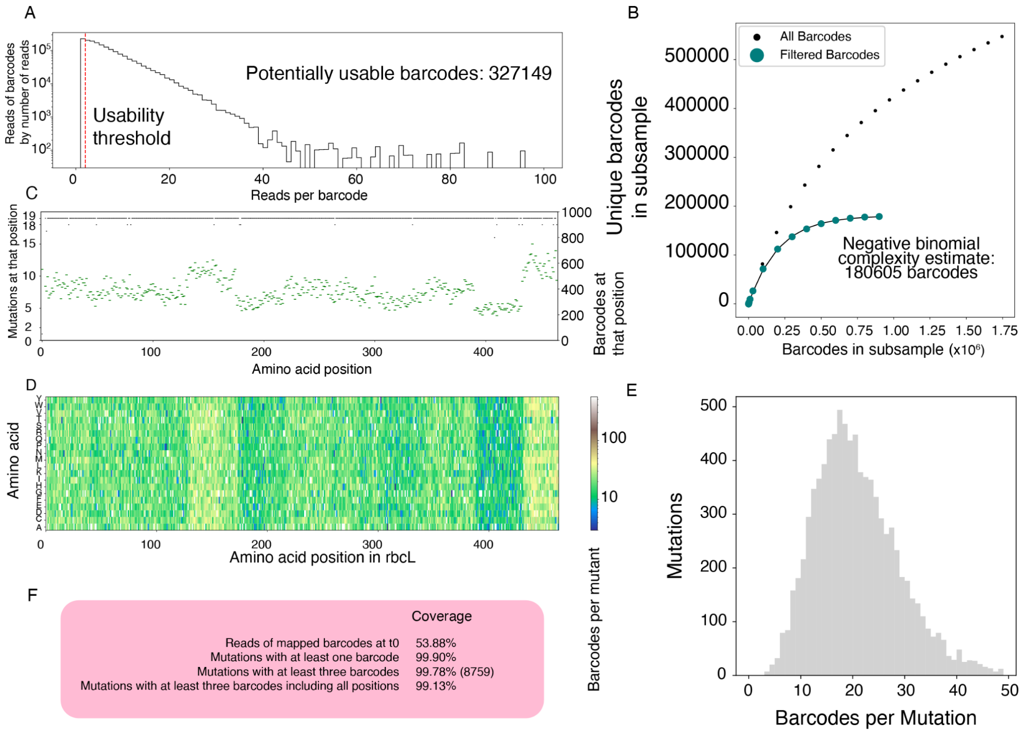


**Figure S6: Library characterization by long-read sequencing. A)** A histogram of reads of plasmids from PacBio sequencing. The y-axis represents the number of reads of plasmids with a given number of reads (i.e. the bar at 50 on the x-axis is as tall as the number of reads of barcodes with 50 reads). We were able to generate a consensus sequence for any barcode with more than 1 read leaving us with 327149 possible barcodes. **B)** A rarefaction plot estimating the overall library complexity, a negative binomial distribution was fit and we estimated a real library complexity of ≈180,000 barcodes. **C)** A plot of how many mutants (of the possible 19) were in our library at each position (black dashes, left axis) and how many barcodes (green dashes, right axis). **D)** A heatmap of how many barcodes were characterized for each mutation. **E)** A histogram of mutants by how many barcodes they had. **F)** Statistics on the completeness of the library. Overall we had >99% of the mutations in our lookup table.

#### *Pairplot at 5% $CO_2$*

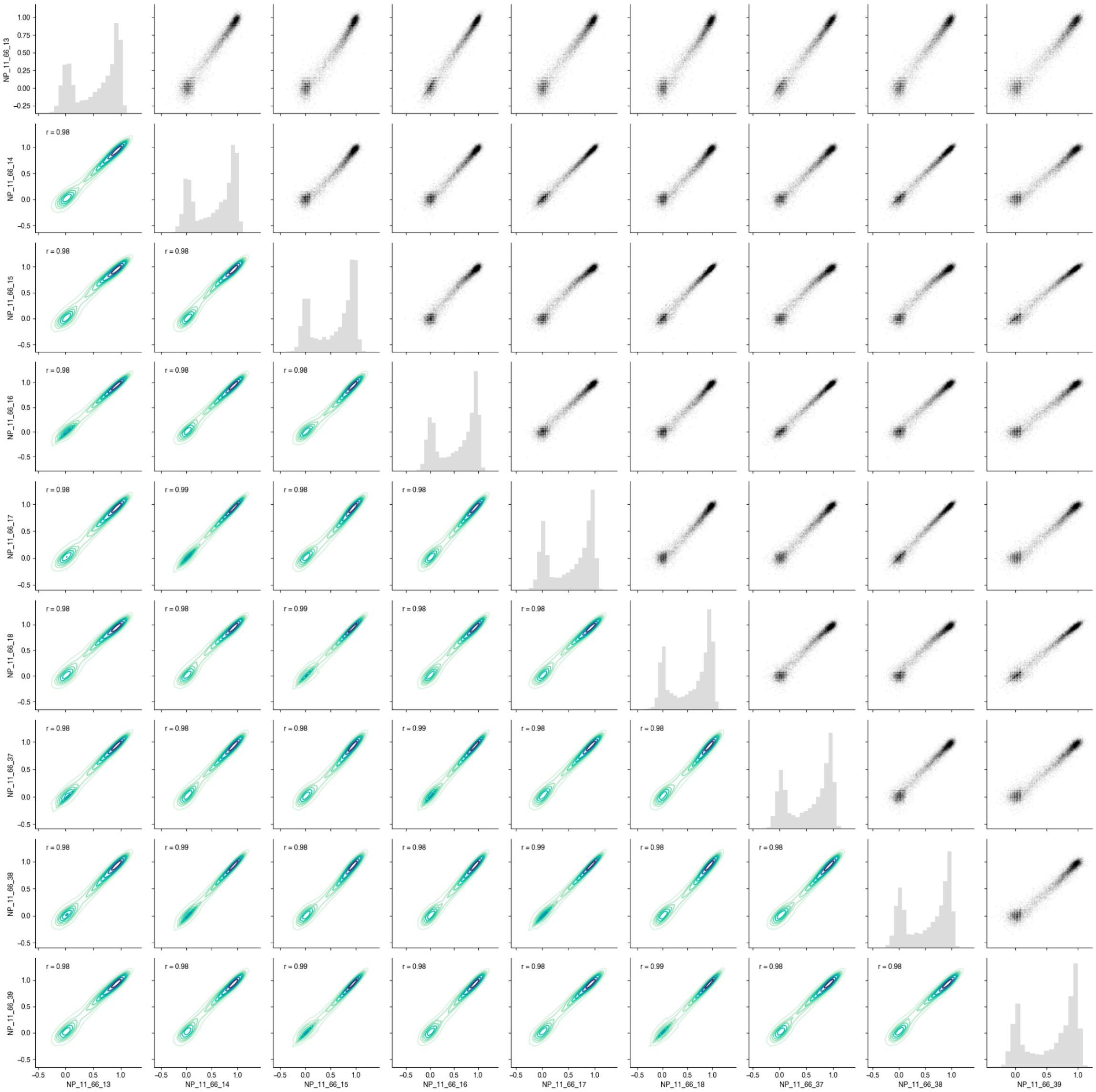


**Figure S7: Pairplots of replicate fitness values:** Fitness values for each mutant are calculated as described in the methods for each replicate individually. These replicates are 3 sets of technical replicates of 3 biological replicates. NP_11_66_13,16 and 37 are technical replicates (same with 14/17/38 and 15/18/39). 37-39 were collected on a different day. Pearson correlations reported for each pair of replicates. The distribution of fitness values is reported along the diagonal and pairwise correlations are reported between replicated off the diagonal. Pearson R is reported in the bottom-left half.

#### *Biochemistry summary*

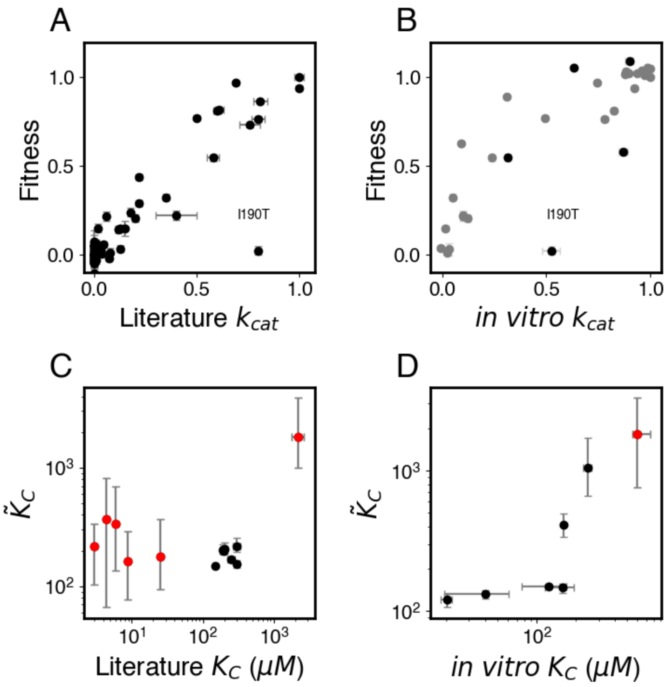


**Figure S8: Comparisons between biochemically measured rubisco kinetic parameters and those same parameters as inferred from fitness values. A and B)** Fitness vs. $k_{cat}$ values, **C and D)** $\widetilde{K}_C$ vs. $K_C$ values. Measurements from the literature in **A** and **C**, values measured in this study in **B** and **D**. Black points in **B** were purified 3 independent times (error bars are standard error), all other data are from individual purifications and have no errors reported. X-axis error bars in A and C are taken from the literature when available. X-axis errors in D and Y-axis errors in A-D are explained in the methods. Outlier mutation is labeled in A and B and is discussed in the supplementary text. Red indicates $\widetilde{K}_C$ estimates with coefficient of variation >1.

##### *Heatmap in pieces*

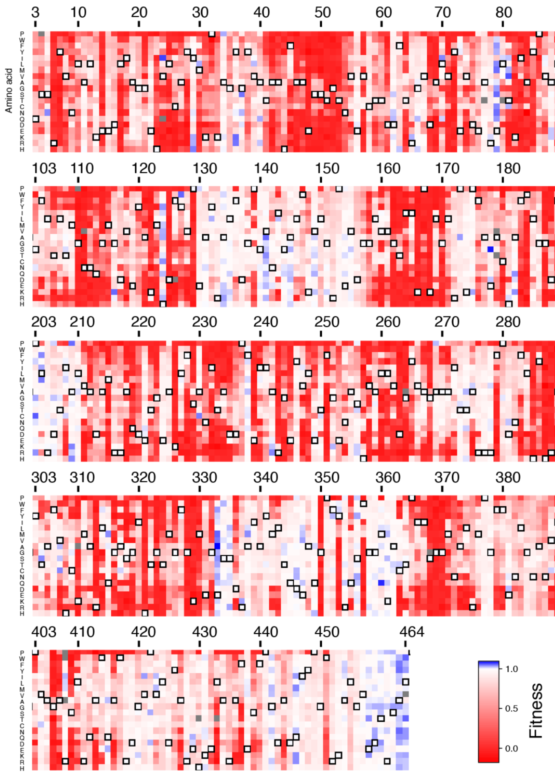


**Figure S9: Full heatmap of fitness values**

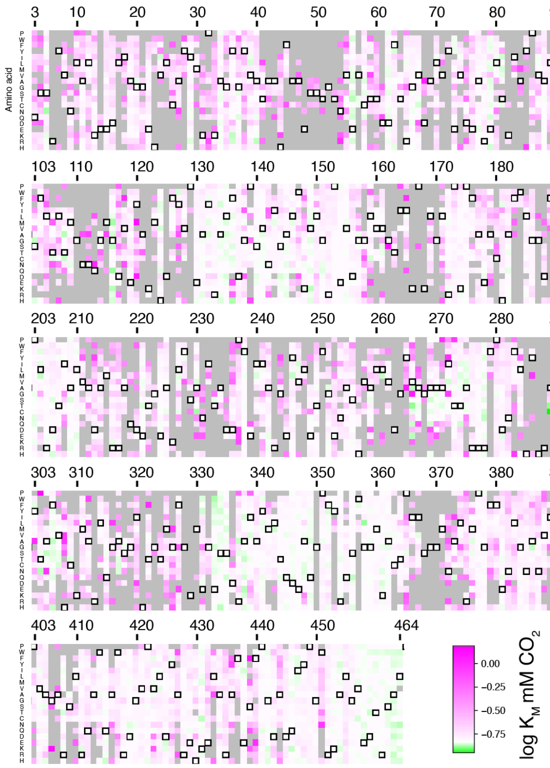


**Figure S10: Full heatmap of $\widetilde{K_C}$ values**

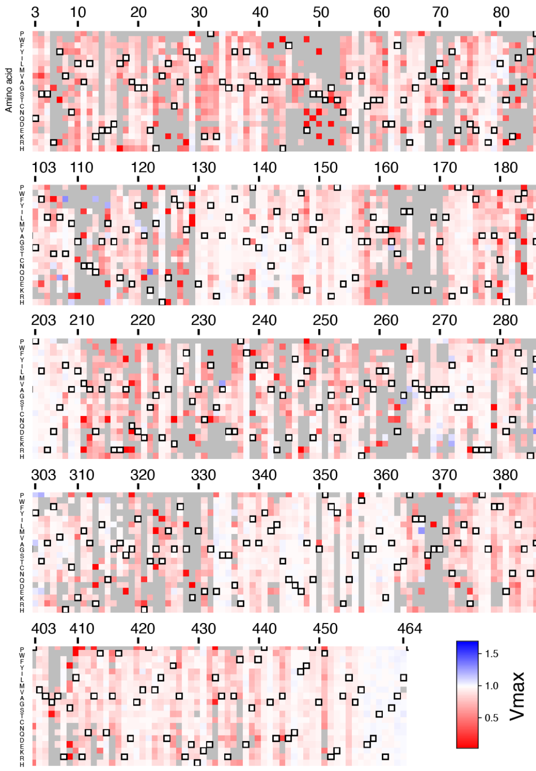


**Figure S11: Full heatmap of $\widetilde{V}_{max}$ values.**

#### *Ridgeline plot*

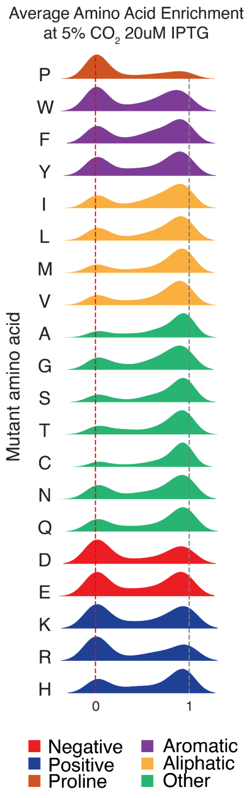

**Figure S12: Histograms of fitness effects of mutations to each amino acid individually.** A histogram of fitness effects of all mutations to the specified amino acid (i.e. the plot for proline is the histogram of the fitness effects of mutations to proline at each position where there isn't a proline naturally). Plots are colored by the type of amino acid.

#### *Conservation vs. Tolerance with 2 different MSAs*

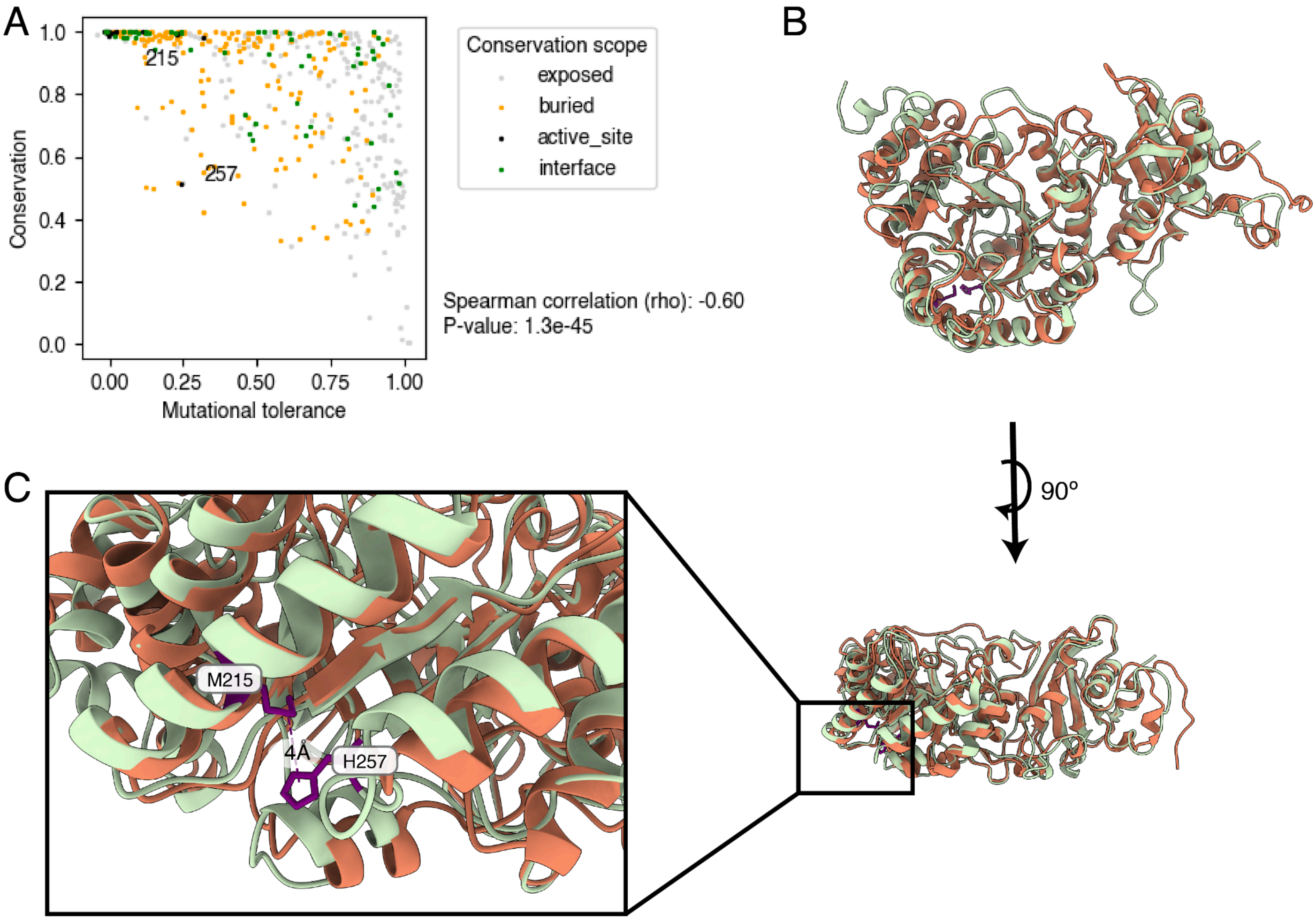


**Figure S13: "Recent" evolution of a tertiary contact: A)** Conservation vs. Tolerance among bacterial Form II rubiscos. As in Figure 2C, mutational tolerance is the average fitness effect of all mutations at a given position. Here conservation is determined from an MSA of all Form II bacterial rubiscos (see methods). Positions 215 and 257 form a tertiary contact in *R. rubrum* and other Form II rubiscos and are thus more conserved than among all rubiscos. **B)** Alignment of 9RUB and 8RUC, *R. rubrum* (green) and spinach (orange) rubisco respectively. **C)** Rotated view and zoom of M215 and H257 from *R. rubrum*. The loop containing them in *R. rubrum* is truncated in spinach.

#### *Pairplots of each other $CO_2$ condition*

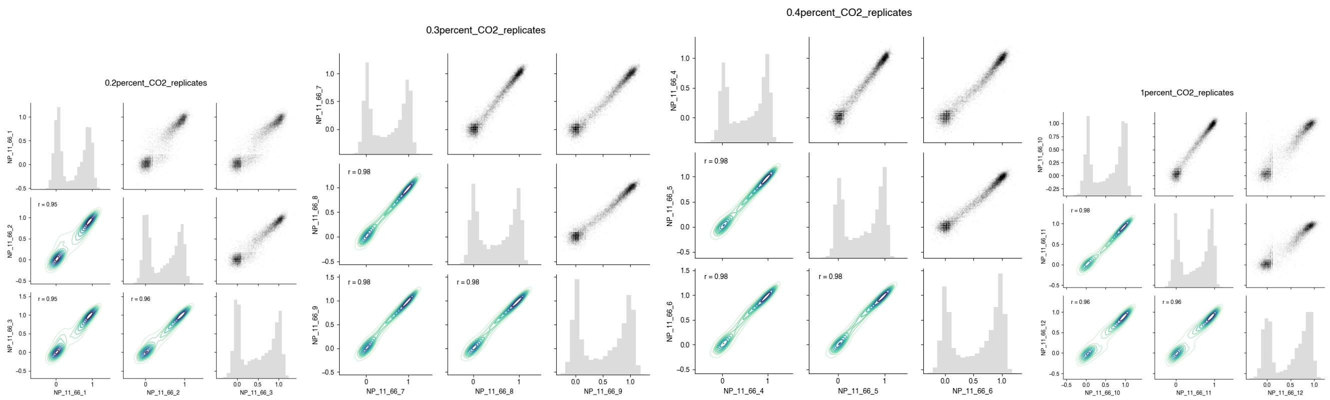


**Figure S14: Pairplots of replicate fitness values at different $CO_2$ concentrations:** Correspondence between three biological replicates of fitness values calculated from the sequenced results of selections at each indicated $CO_2$ concentration. NP_11_66_1/2/3 are at 0.2% $CO_2$, 7/8/9 are at 0.3%, 4/5/6 are at 0.4% and 10/11/12 are at 1%. The distribution of fitness values is reported along the diagonal and pairwise correlations are reported between replicated off the diagonal. Pearson R is reported in the bottom-left half.

*Pairplots between conditions*

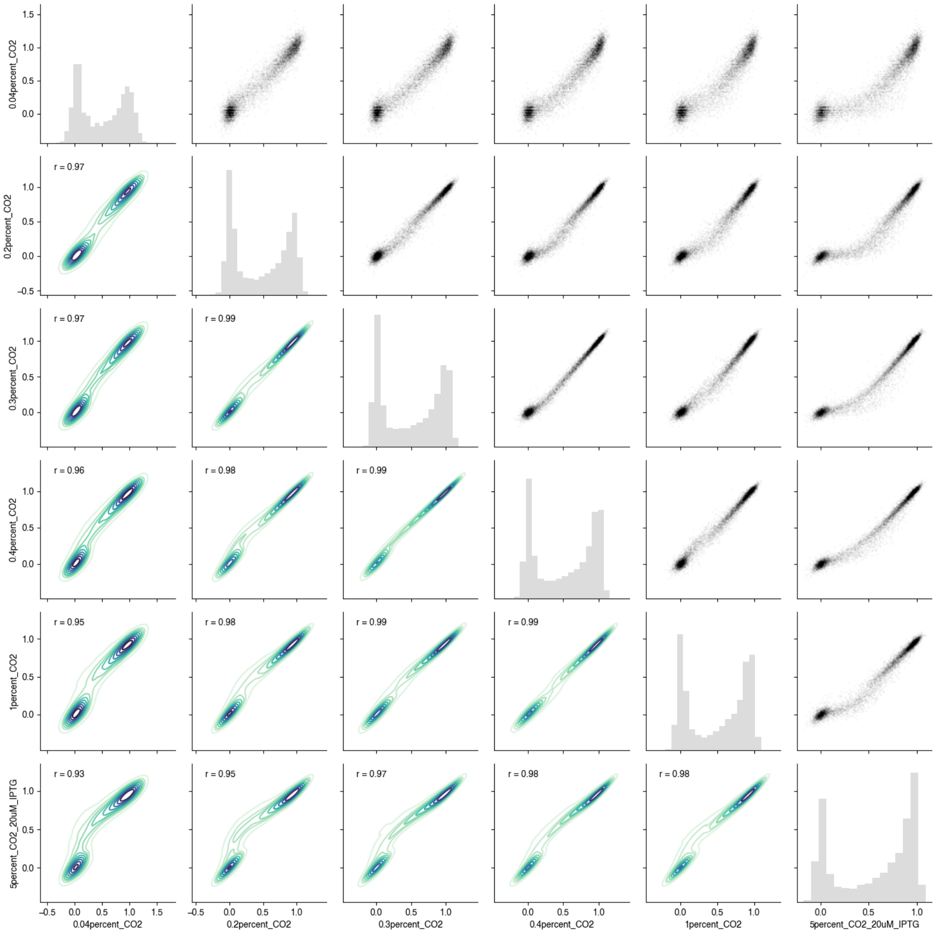


**Figure S15: Pairplots of fitness values at different $CO_2$ concentrations:** Correspondence between fitness values at each $CO_2$ concentration. The distribution of fitness values is reported along the diagonal and pairwise correlations are reported between replicated off the diagonal. Pearson R is reported in the bottom-left half. As shown in figure 3 the distribution shifts to higher fitness values at higher $CO_2$ concentrations. The two peaks of the bimodal distribution also become more distinct with increasing $CO_2$ indicating that the difference between functional and non-functional variants is more extreme, likely reflecting the strength of selection.

*$\widetilde{V}_{max}$ vs. Fitness*

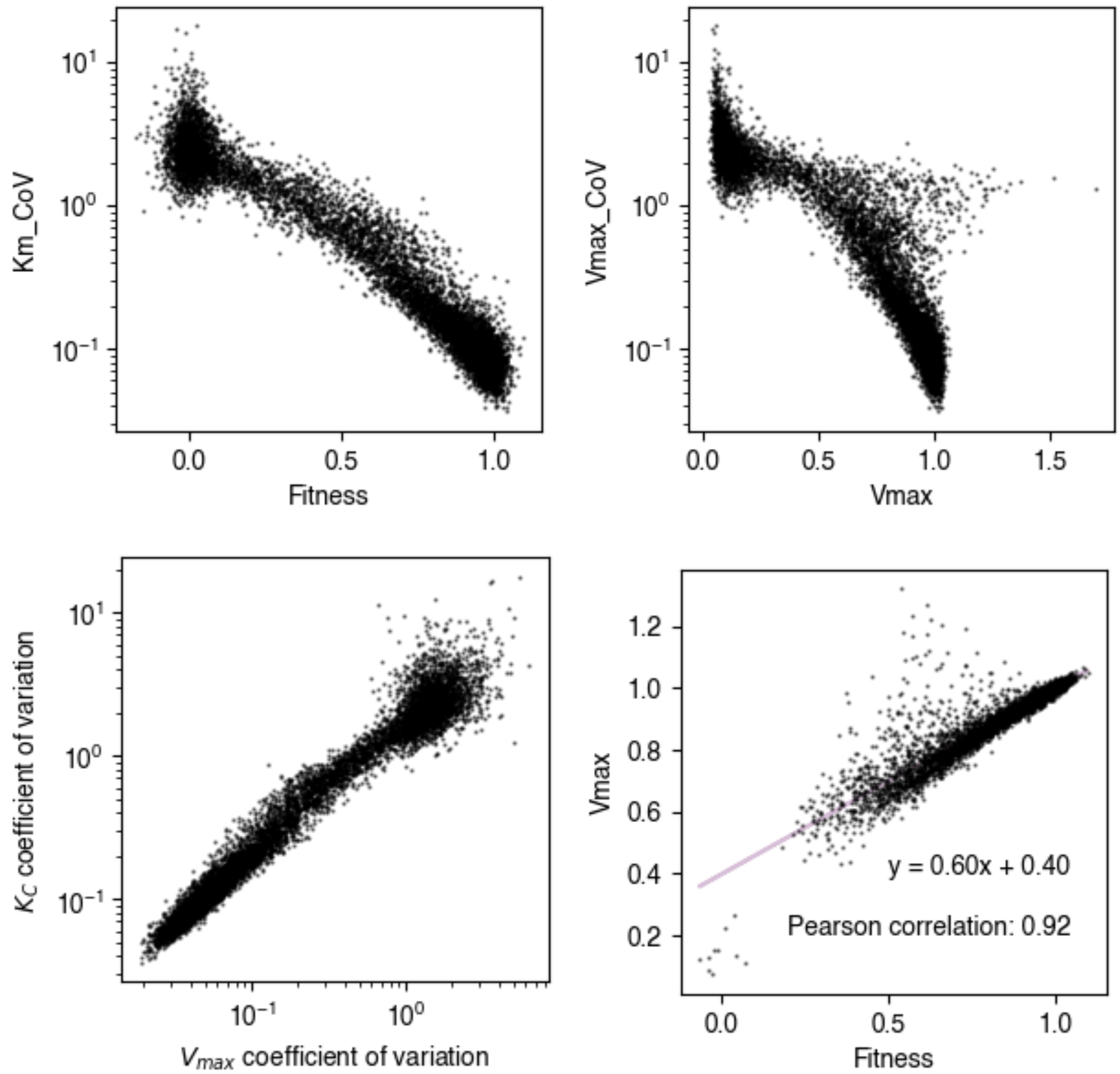


**Fig. S16: Correlations of CoV, fitness and $\widetilde{V}_{max}$: A)** $\widetilde{K}_C$ coefficient of variation as a function of fitness. **B)** $\widetilde{V}_{max}$ coefficient of variation as a function of $\widetilde{V}_{max}$ **C)** $\widetilde{K}_C$ coefficient of variation as a function of fitness $\widetilde{V}_{max}$ coefficient of variation. **D)** Correlation of $\widetilde{V}_{max}$ and Fitness. Only mutants with a coefficient of variation <1 are plotted here; those mutants typically have low fitness and are thus harder to fit to a Michaelis-Menten model.

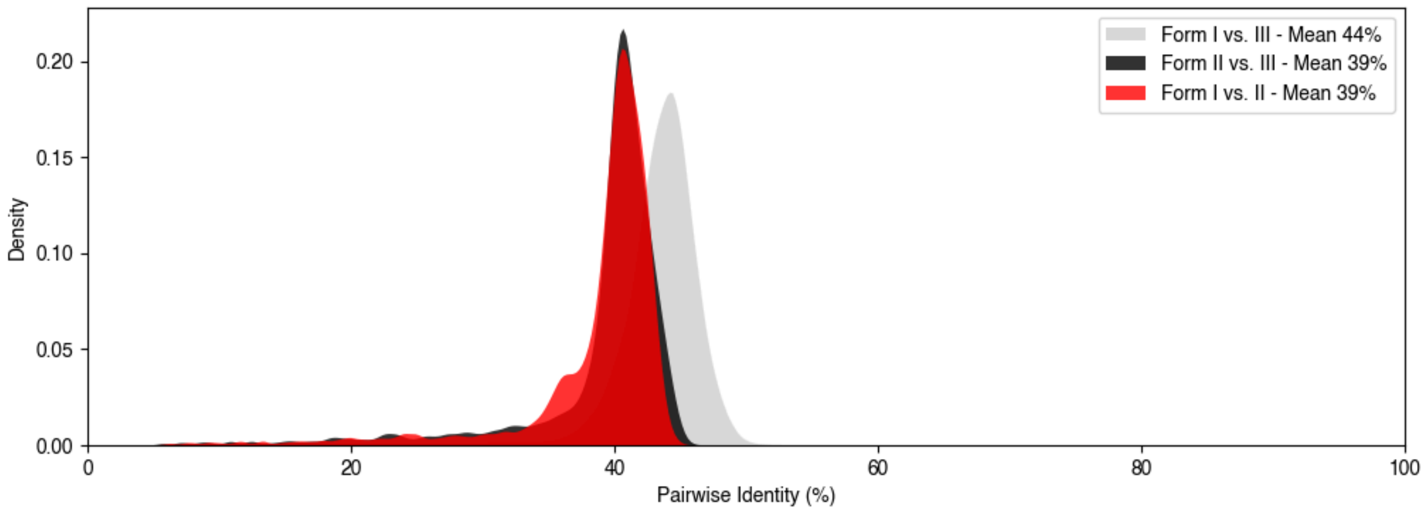


**Figure S17: Pairwise identities between rubisco sequences across Forms:** Representative rubisco sequences from (*8*) were compared for pairwise identity. Form I sequences were picked to have a maximum sequence identity between one another of 85% in order to sample sequences more evenly (out of fear of oversampling plant sequences). Form II and III sequences were chosen randomly.

#### Tables and files

##### *Table S1*

| | $k_{cat}$ ($s^{-1}$) | $K_M$ (µM $CO_2$) |
|---|---|---|
| WT | 9.9 ± 0.4 | 148 ± 10 |
| V266T | 5.1 ± 0.1 | 87 ± 5 |
| A102Y | 1.7 ± 0.04 | 53 ± 3 |

##### *Data File S1*

Literature and in vitro kinetics data

##### *Data File S2*

Enrichments, $\widetilde{V}_{max}$, $\widetilde{K}_C$ and associated errors.

#### Plasmids and Primers

##### *Data file S3*

**Plasmids:**
NP-11-64-1
pET28_SUMO

**Primers:**
Illumina sequencing
Mutagenic primers for $K_M$18 library
Library prep primers (hunks)

TWIST oligos

#### Discussion of outlier in Figure 1E

I190T was the only outlier in our comparison of *in vitro* $k_{cat}$ measurements from the literature and our fitness data. Because the value was reported without error estimates (*32*), we re-measured the $k_{cat}$ of this mutant and found it to be 4.24 $s^{-1}$, which is 52% of the WT value, down from 80% previously reported. Still, the value appears to be anomalous compared to the rest of the trend (Fig. S8B). One potential explanation is that the mutation at that position has a strong negative effect on protein expression. Another possibility, given that I190T is adjacent to the key active site lysine, K191, is that I190T causes a negative effect on lysine carbamylation that is, for some reason, more pronounced *in vivo* than *in vitro*. It is hard to explain why that would be the case because the $CO_2$ concentrations are the same.

#### Derivation of Michaelis-Menten Fit

Following Stiffler et al. we assume that the differences in bacterial growth rate are proportional to the differences in growth-limiting enzymatic activity.

(Eq S1) $$\mu_{mut} - \mu_{wt} \propto v^{ru}_{mut} - v^{ru}_{wt}$$

Under the presumption of log-phase growth, the expected log ratio of reads after elapsed time *t* and normalized to the wild-type reads is given by,[1]

(Eq S2) $$e_{mut} = log_{10}\left(\frac{N_{mut,f}}{N_{mut,0}}\right) - log_{10}\left(\frac{N_{wt,f}}{N_{wt,0}}\right)$$

Substituting in the condition of exponential growth, ie. $N_{i,f} = N_{i,0}\, e^{\mu_i t}$, and simplifying yields,

(Eq S3) $$e_{mut} = \frac{t}{ln\,10}\left(\mu_{mut} - \mu_{wt}\right)$$

In order to normalize the enrichments, we divide by the log enrichment of the wild-type counts relative to the median enrichment of variants with mutated catalytic residues (and thus catalytically dead rubisco). We then add one for the convention that dead variants be centered at an enrichment of zero and that wild-type be at an enrichment of one. Thus, the normalized mutant enrichment is,

(Eq S4) $$e_{mut,\,norm} = \frac{log_{10}\left(\frac{N_{mut,f}}{N_{mut,0}}\right) - log_{10}\left(\frac{N_{wt,f}}{N_{wt,0}}\right)}{log_{10}\left(\frac{N_{wt,f}}{N_{wt,0}}\right) - \langle\, log_{10}\left(\frac{N_{dead,f}}{N_{dead,0}}\right)\rangle} + 1$$

Then substituting Eq S3 we obtain,

(Eq S5) $$e_{mut,\,norm} = \frac{\mu_{mut} - \mu_{wt}}{\mu_{wt} - \mu_{dead}} + 1$$

Using the assumption in Eq S1 and the fact that the enzyme velocity of dead mutants is zero we obtain the expected normalized enrichment as a function of the rubisco velocities,

(Eq S6) $$e_{mut,\,norm} = \frac{v_{mut}}{v_{wt}}$$

[1]Note that Eq 2 would also contain a normalization factor to account for the total number of reads obtained for the pre- and post-selection conditions. It is, however, a common factor for both the mutant and wild-type counts and therefore cancels out. Furthermore, the real analysis also includes pseudo-counts which are omitted here in the derivation of the fit equation for simplicity.

Finally, using the Michaelis-Menten equation we obtain the predicted enrichments as a function of $CO_2$ concentration and the enzyme kinetic parameters.

(Eq S7)
$$e_{mut,\, norm}\left([CO_2]\right) = \frac{V_{max,\, mut}\left(K_{M,wt}+[CO_2]\right)}{V_{max,wt}\left(K_{M,mut}+[CO_2]\right)}$$

Thus, in practice, we use Eq S7 as the fit equation to the normalized enrichment values for each variant across a range of $CO_2$ concentrations. For each we have as fit parameters the ratio of maximum velocities between the mutant and wild-type, $V_{max,mut} / V_{max,wt}$, and the mutant $K_C$ with the wild-type $K_C$ set to the literature value of 149 μM.
